# ADP-ribosylation of mitochondrial proteins is mediated by Neuralized-like protein 4 (NEURL4)

**DOI:** 10.1101/2020.12.28.424513

**Authors:** Maria Dafne Cardamone, Yuan Gao, Julian Kwan, Vanessa Hayashi, Megan Sheeran, Junxiang Xu, Justin English, Joseph Orofino, Andrew Emili, Valentina Perissi

**Affiliations:** Department of Biochemistry, Boston University School of Medicine, Boston, MA 02118, USA; Center for Network Systems Biology, Boston University, Boston, MA 02118, USA

## Abstract

ADP-ribosylation is a reversible post-translational modification where an ADP-ribose moiety is covalently attached to amino acid side-chains of target proteins either as mono-ADP-ribose (MARylation or MAR) or poly-ADP-ribose chains (PARylation or PAR) by a class of enzymes called ADP-ribosyltransferases (ARTs). Although ADP-ribosylation is best known for its nuclear roles, ADP-ribosylation of extra nuclear proteins is increasingly recognized as a key regulatory strategy across cellular compartments. ADP-ribosylation of mitochondrial proteins, in particular, has been widely reported, even though the extent to which ADP-ribosylation of specific proteins regulates mitochondrial functions is unclear and the exact nature of mitochondrial ART enzymes is debated.

Here, we have identified Neuralized-like protein 4 (NEURL4) as a mitochondrial ART enzyme and profiled the NEURL4-dependent ADP-ribosylome in mitochondrial extracts from Hela cells by LC-MS/MS, using isobaric tandem mass tag (TMT) labeling for relative quantification. Comparison of WT and NEURL4-KO cells generated by CRISPR/Cas9 genome editing revealed that most ART activity associated with mitochondria is lost in absence of NEURL4. Putative NEURL4 targets include numerous mitochondrial proteins previously shown to be ADP-ribosylated. In particular, we show that NEURL4 enzymatic activity is required for the regulation of mtDNA integrity via poly-ADP-ribosylation of mitochondrial specific Ligase III (mtLIG3), the rate-limiting enzyme for mitochondrial DNA (mtDNA) Base Excision Repair (BER).

Collectively, our studies reveal that NEURL4 acts as the main mitochondrial ART enzyme under physiological conditions and provide novel insights in the regulation of mitochondria homeostasis through ADP-ribosylation.

## Introduction

Protein ADP-ribosylation is a widespread post-translational modification catalyzed by members of a family of enzymes known as ADP-ribosyltransferases (ARTs). ARTs covalently link ADP-ribose moieties, either as mono-ADP-ribose (MARylation or MAR) or poly-ADP-ribose chains (PARylation or PAR), on target proteins using NAD^+^ as cofactor (Aravind et al., 2015; Cohen and Chang, 2018; Kraus, 2015). ADP-ribosylation can occur on a number of amino acid residues, including serine, glutamate, aspartate, arginine, lysine and cysteine(Altmeyer et al., 2009; Barkauskaite et al., 2015; Bonfiglio et al., 2017; Leslie Pedrioli et al., 2018; McDonald and Moss, 1994; Messner and Hottiger, 2011; van Ness et al., 1980; Palazzo et al., 2018; Vyas et al., 2014; Welsh et al., 1994). ARTs are classified in two major subclasses depending on their conserved structural features: ARTCs (cholera toxin-like) and ARTDs (diphtheria toxin-like). The eukaryotic ARTDs, most commonly known as poly-(ADP-ribose) polymerases (PARPs), form the largest and most characterized group of ARTs, composed of 17 members, with PARP1 (ARTD1) being the founding member (Amé et al., 2004; Hottiger et al., 2010). Although the overall structure of ARTD catalytic domain is highly conserved, differences in the critical amino acidic triade H-Y-E at the active center of the catalytic domain determine the specific enzymatic ability of each member of the family (Aravind et al., 2015; Cohen and Chang, 2018; Hassler and Ladurner, 2012; Otto et al., 2005). In accord with the H and Y residues being required for binding to NAD^+^ and the E playing a key role in the elongation process, members of the ARTD family capable to catalyze PARylation, including PARP1, present the full H-Y-E motif, whereas enzymes performing MARylation contain a modified H-Y-I motif. Catalytically inactive enzymes that are unable to efficiently bind NAD^+^ lack the H (Aravind et al., 2015; Cohen and Chang, 2018; Hottiger et al., 2010).

In mammals, ADP-ribosylation is best known for its role in mediating the DNA damage response in the nucleus (Azarm and Smith, 2020; Gupte et al., 2017; Ray Chaudhuri and Nussenzweig, 2017). However, in the past two decades, a growing literature has provided evidence that ADP-ribosylation of proteins across all cellular compartments is relevant to a wide range of biological functions, including metabolism, inflammation, cancer and aging (Bai and Cantó, 2012; Fehr et al., 2020; Kraus, 2015; Leung, 2014; Slade, 2020; Szántó and Bai, 2020; Vyas et al., 2013). While ARTs in the nucleus and cytoplasm have been extensively documented, the presence of ADP-ribosylation enzyme/s in the mitochondria remain controversial (Brunyanszki et al., 2016; Masmoudi and Mandel, 1989). Although ART activity in mitochondria extracts from different tissues has been reported for decades (Burzio et al., 1981; Kun et al., 1975; Masmoudi and Mandel, 1987; Masmoudi et al., 1988; Richter et al., 1983), and despite widespread reports of ADP-ribosylated mitochondrial proteins (Herrero-Yraola et al., 2001; Innovation and Hendriks, 2019; Martello et al., 2016; Pankotai et al., 2009; Vivelo et al., 2017; Williams et al., 2018), the specific nature of a mitochondrial ART has remained elusive. Among others, ARTD1/PARP1 has been investigated as a mediator of ADP-ribosylation in mitochondria based to its ability to interact with Mitofillin (Rossi et al., 2009). Based on PARP1 best known role as a regulator of the genomic DNA (gDNA) damage repair pathway in the nucleus, it was proposed that intramitochondrial PARP1 regulates mtDNA stability by interacting with DNA ligase III (LIG3) on mtDNA (Brunyanszki et al., 2016; Rossi et al., 2009). However, there is no direct evidence that PARP1 enzymatic activity is required for the regulation of mtDNA maintenance nor that PARP1 can mediate the PARylation of proteins involved in the DNA damage response or that of other targets identified as part of the mitochondrial PARylome. Moreover, it was recently shown that, in nonstressed Hela cells, PARP1/ARTD1 inhibition by Olaparib eliminated nuclear parylation, without affecting extra-nuclear ADP-ribosylation signal (Nowak et al., 2020). This indicates that, even though PARP1/ARTD1 is constitutively activated under physiological conditions, its activity only accounts for nuclear modifications, thus raising the outstanding question of whether another ARTD family member or a previously unknown ART enzyme is responsible for mitochondrial PARylation.

*In silico* analyses aimed at identifying novel NAD^+^-utilizing enzymes recently predicted the presence of an H-Y-E like ART domain in the C-terminal domain of Neuralized-like protein 4 (NEURL4) (de Souza and Aravind, 2012), indicating that NEURL4 may be an additional member of the ARTD family. NEURL4 is currently classified as a member of the Neuralized family based on the presence of six Neuralized Homology Repeat (NHR) domains. The NHR domain is highly conserved from flies to mammals and it is important in the mediation of protein-protein interactions. Four Neuralized family members have been identified in mammals to date. NEURL1, NEURL2 and NEURL3 contain either a RING or SOCS domain, in addition to the NHR domains, and therefore serve as E3 ubiquitin ligases in a variety of physiological processes(Liu and Boulianne, 2017). NEURL4 and its fly homologue, *d*NEURL*4* lack of other recognizable domains and have not been associated with intrinsic ubiquitination activity, even though it is appreciated that NEURL4 promotes ubiquitin signaling through interaction with the HECT E3 ubiquitin ligase HERC2 (Al-Hakim et al., 2012; Jones and Macdonald, 2015; Li et al., 2012; Loukil et al., 2017). Together, NEURL4 and HERC2 localize to the centrosome and interact with CP110 to regulate centrosome morphology and centriolar homeostasis via CP110 ubiquitin-dependent degradation (Al-Hakim et al., 2012). NEURL4/HERC2 complexes also regulate MDM2-p53 dimerization in response to DNA damage and Notch signaling via Delta-like 1 (Dll1)/Delta (Dl) recycling, even though in both cases HERC2 enzymatic activity appeared dispensable (Cubillos-Rojas et al., 2017; Imai et al., 2015).

Here, we report evidences that NEURL4 serves as a mitochondrial ART and discuss how the characterization of NEURL4 role in mediating the PARylation of a large number of mitochondrial proteins has led to novel insights into the relevance of ADP-ribosylation for the maintenance of mitochondrial homeostasis.

## Results

Comparative structure and genomic analysis aimed at uncovering novel NAD^+^-utilizing enzymes have predicted the existence of a putative H-Y-E[D-Q] ADP-ribosyltransferase domain (ART), similar to that of ARTD1, within the C-terminal domain of NEURL4 (de Souza and Aravind, 2012)(**Fig. 1A**). Based on this prediction and previous reports of NEURL4 association with centromeres-associated factors (Al-Hakim et al., 2012; Loukil et al., 2017), it was suggested that NEURL4 might be regulating centrosomal assembly through poly-ADP-ribose polymerase (PARP)-like activity(Aravind et al., 2015; de Souza and Aravind, 2012). However, our profiling of its subcellular localization by immunohistochemistry revealed that in Hela cells NEURL4 was predominantly found in the mitochondria (**Fig. 1B**). In particular, the pattern of digestion by Proteinase K, as compared to that of known markers of different mitochondrial compartments, indicated that NEURL4 localized to the mitochondrial matrix (**Fig. 1C**). This was confirmed by electron microscopy, with immunogold labelling of NEURL4 showing its proximity to the cristae of the inner mitochondrial membrane (IMM) (**Fig. 1D**). Based on the identification using the Mitoprot software (Claros, 1995) of a putative mitochondrial targeting sequencing (MTS) within aa 1 to 65 of NEURL4 (see **Fig. 1A**), we predicted that the import of NEURL4 into the mitochondria would occur through the classic import pathway with cleavage of the N’terminal presequence. Processing was confirmed in transiently transfected Hela cells by the accumulation of uncleaved NEURL4, as visualized by WB for the N-terminal Flag tag, upon transient knockdown of mitochondrial processing peptidases (MPP) alpha and beta by siRNAs (**Fig. 1E)**.

**Figure 1.**
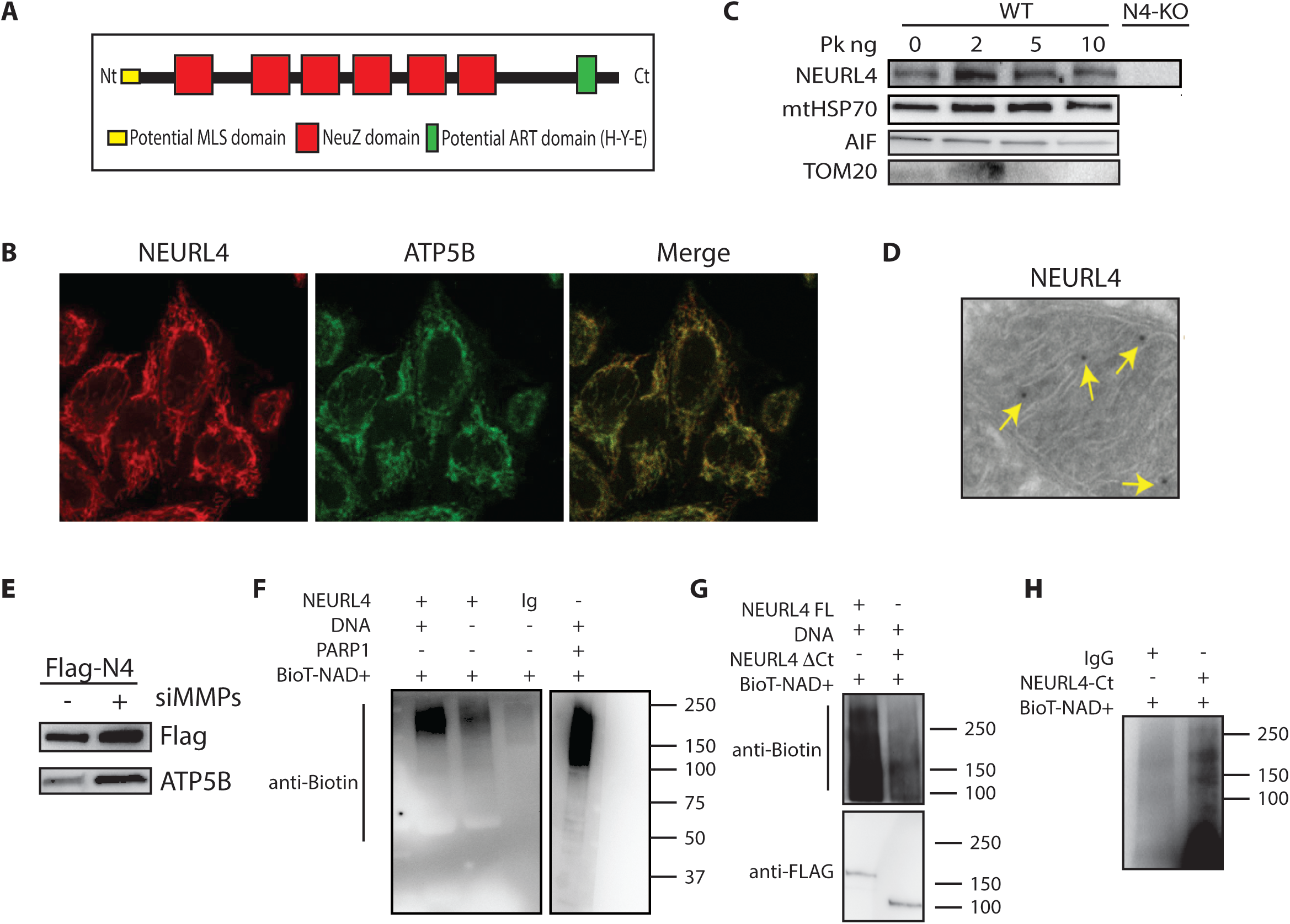
NEURL4 is a mitochondrial protein with a functional ART domain. **(A)** Graphical representation of NEURL4 known and predicted structural domains. **(B)** NEURL4 localization to the mitochondria by IHF staining in Hela cells as shown by costaining with the mitochondrial marker ATP5B. **(C)** Proteinase K protection assay showing NEURL4 localization to the mitochondrial matrix. Degradation patterns of TOM20, AIF1 and mtHSP70 are representative of proteins respectively in the OMM, IMM and matrix. **(D)** Mitochondria labeling by Immunogold EM for NEURL4 in Hela cells. **(E)** Increased FLAG-NEURL4 stability upon down regulation of MMPs (both MMPa and MMPβ) by specific siRNA. **(F-G-H)** *In vitro* PARylation assays showing NEURL4 ability to produce poly-ADP-ribosylation chains (**F**) via its Ct domain (**G** and **H**). Immuno-purified endogenous NEURL4, FLAG-NEURL4-ΔCt or FLAG-NEURL4-Ct was incubated with biotin-NAD^+^ in presence or in absence of activated DNA. Recombinant PARP1/ARTD1 was used as positive control (**F**).

These results indicated that NEURL4 was a previously unrecognized mitochondrial protein and suggested it was a promising candidate for mediating ADP-ribosylation in mitochondria. To confirm the predicted enzymatic activity, we performed an *in vitro* PARylation assays with NEURL4 isolated from Hela extracts, including PARP1/ARTD1 as positive control. As shown in **Figure 1F**, NEURL4 mediated the synthesis of poly-ADP-ribose (PAR) chains *in vitro,* with its activity being potentiated in presence of activated DNA (**Fig. 1F lane 1**). A truncated form of NEURL4, lacking the predicted enzymatic domain at the C-terminus, failed to generate PAR chains under the same conditions (**Fig. 1G**), while the NEURL4 C-terminus alone was sufficient to record ADP-ribosylation activity in mitochondrial extracts (**Fig. 1H**).

To further investigate the relevance of NEURL4 enzymatic activity and characterize its role in the mitochondria, we used CRISPR/Cas9 genome editing in Hela cells to generate two independent NEURL4 knock-out (N4-KO1 and N4-KO2) cell lines (**Fig. 2A** and **2B**). In both models, we observed a dramatic reduction of mitochondria-associated ADP-ribosylation in absence of NEURL4, as monitored by western blotting of mitochondrial extracts with anti-PAR antibody (**Fig. 2C**). This phenotype was rescued by re-introducing in N4-KO cells either the full length NEURL4 or the catalytic domain alone (**Fig. 2D**), indicating that NEURL4 enzymatic activity is required for mediating mitochondrial PARylation. Moreover, the mitochondrial PARylation observed upon reintroducing NEURL4 was impaired by treatment with BGP-15 2HCl, a generic inhibitor of PAR activity that would impact upon NEURL4 activity, but not in presence of an inhibitor specific for PARP1/ARTD1 (Olaparib) which instead appeared to potentiate PARylation (**Fig. 2E**), thus further excluding the possibility of PARP1/ARTD1 being involved in NEURL4-mediated activity. Lastly, we assessed the acetylation status of mitochondrial proteins. Because the mitochondrial pool of NAD^+^ is kept segregated from that of cytosol/nucleus and NAD^+^ is a shared cofactor of ART/SIRT enzymes, we reasoned that, if NEURL4 was a major user of NAD+ for ADP-ribosylation, in its absence the increased availability of mitochondrial NAD^+^ would favor the activity of other enzymes that require NAD^+^ as cofactor, such as NAD-dependent deacetylases. Indeed, the acetylation level of mitochondrial proteins was significantly decreased in N4-KO cells, as expected in presence of elevated mitochondrial sirtuins activity, while no changes were observed in the abundance of nuclear acetylation (**Fig. 2F**).

**Figure 2.**
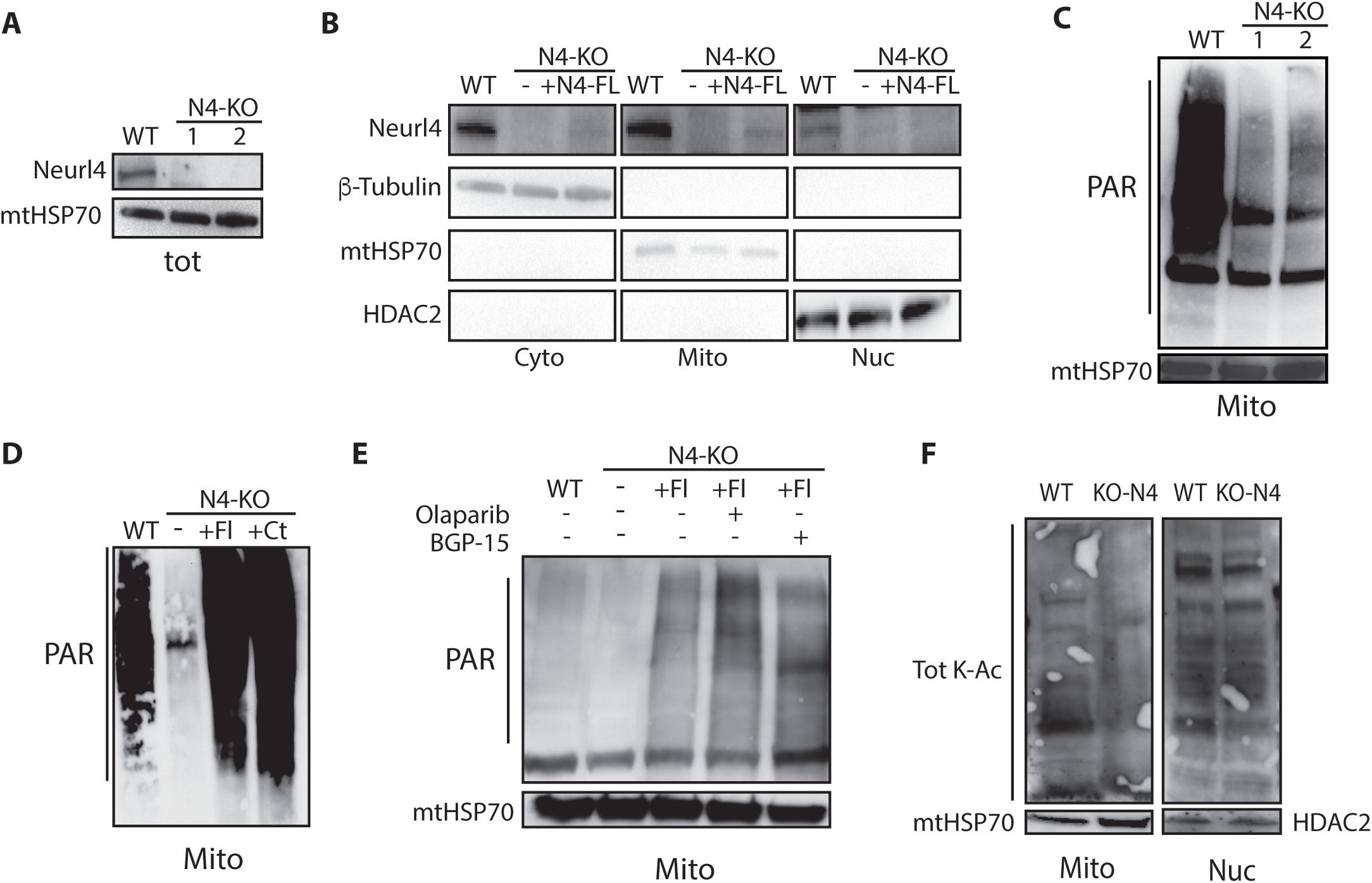
NEURL4 enzymatic activity is essential for mitochondrial PARylation. **(A and B**) Validation of NEURL4 KO cell lines generated by CRISPR/Cas9 genome editing of Hela cell using two independent gRNA. WB of whole cell extracts **(A)** or fractionated cytosolic, mitochondrial and nuclear extracts **(B)**. Efficiency of sub-cellular fractionation is confirmed by WB for Beta-tubulin, mtHSP70 and HDAC2). **(C)** WB anti-PAR on mitochondrial protein extracts showing that most mitochondrial PAR production is lost in untreated Hela cells upon NEURL4 deletion. **(D)** WB anti-PAR on mitochondrial protein extracts from Hela cells showing that loss of ADP-ribosylation in absence of NEURL4 can be rescued reintroducing either full length NEURL4 or the Ct domain alone. **(E)** NEURL4 enzymatic activity is inhibited by the generic PARP inhibitor BGP-15 2 HCl (10μg/ml) but not by PARP1/ARTD1 inhibitor Olaparib (1μM). **(F)** WB anti-Acetylated Lysine on mitochondrial and nuclear extracts from WT and N4-KO cells.

Next, we asked whether the newly identified NEURL4 enzymatic activity was required for maintaining mtDNA integrity. Poly-ADP ribosylation is well established as a key regulatory process for the repair of damaged nuclear DNA (nDNA)(Azarm and Smith, 2020) and previous studies indicate it is likely to play a similar role for mtDNA, even though the identity of the responsible enzyme is unclear (Brunyanszki et al., 2016; Kadam et al., 2020; Szczesny et al., 2014). To assess NEURL4 role in this process, we first confirmed that it is found associated with mtDNA by Chromatin Immunoprecipitation (ChIP) (**Fig. 3A**). Then, we assessed the amount of Poly-ADP-Ribosylation associated with mtDNA in presence or absence of NEURL4 and observed a significant decrease in N4-KO cells in comparison to their parental line (**Fig. 3B**). Together, these findings indicate that enzymatically active NEURL4 is present the mtDNA. Because N4-KO cells have a comparable mitochondria content than their parental line (**Fig. 3C**), it is unlikely that the loss of NEURL4 has any significant effect on DNA replication. However, by long-range PCR we observed a significantly higher amount of mtDNA deletions in N4-KO cells as compared to wild type (**Fig. 3D**). These findings, together with the previous observation of NEURL4 enzymatic activity being enhanced in presence of activated DNA (see **Fig. 1E**), as characteristic of nuclear PARP1/ARTD1, are supportive of NEURL4 contributing to the regulation of mtDNA damage repair.

**Figure 3.**
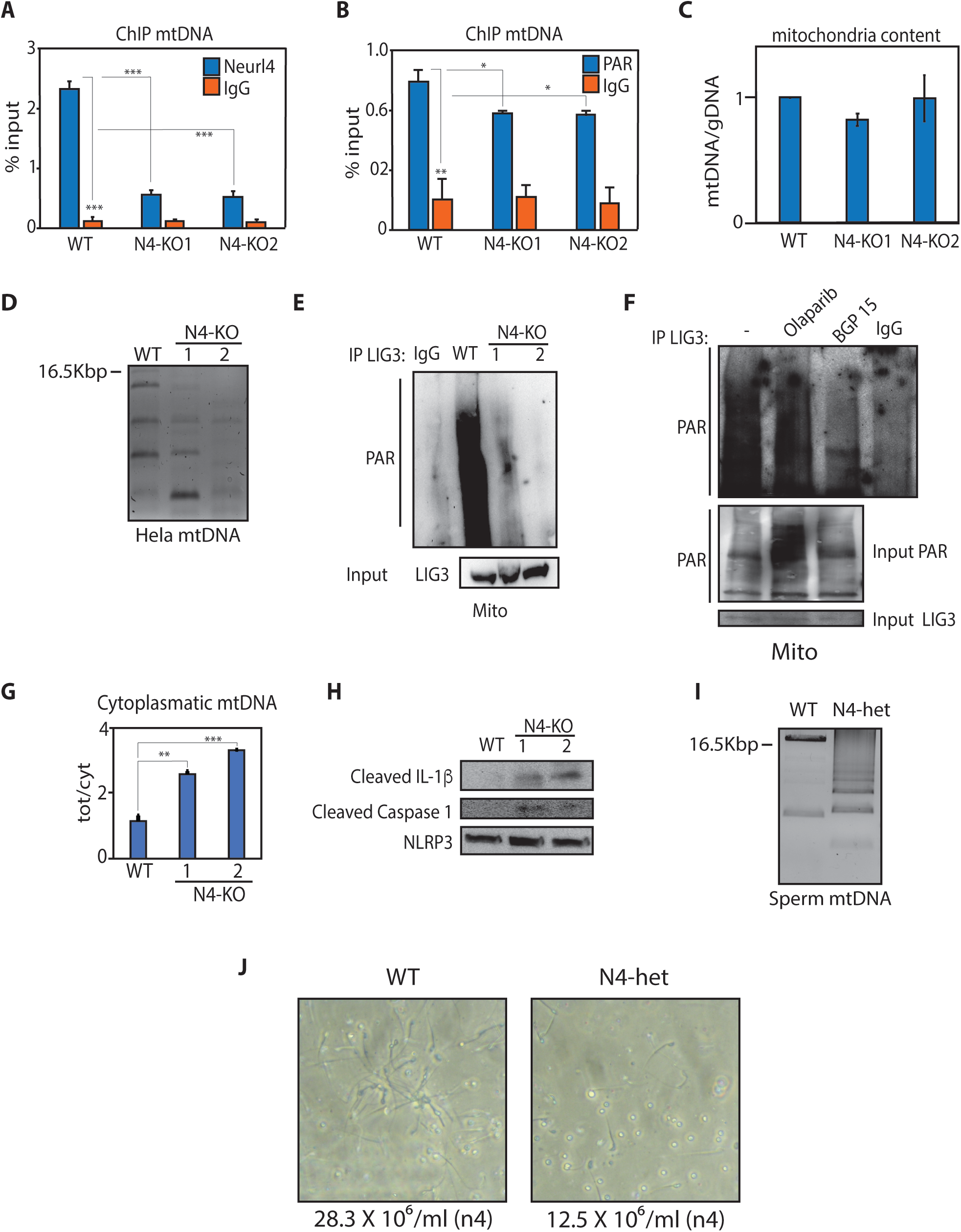
NEURL4 is required for the regulation of mtDNA integrity. **(A and B**) ChIP analysis showing recruitment of NEURL4 at mtDNA sites (**A**) and decreased levels of Poly-ADP-Ribosylation associated with mtDNA in N4-KO cells (**B**). Plots include representative results obtained in three distinct biological replicate experiments, with bar graphs representing the mean +/− standard deviation (SD) between technical triplicates of one single experiment. **(C)** No significant change in mitochondrial content measured by PCR ratio between mtDNA and nuclear DNA. **(D)** Long range PCR assay showing increased mtDNA deletions in N4-KO cells as compared to the Hela parental line (WT). **(E and F)** Immunoprecipitation assay showing suppression of mtLIG3 parylation in absence of NEURL4 (**E**) or in presence of the generic PARP inhibitor BGP-15 2 HCl (**F**). **(G)** qPCR assay showing increased mtDNA accumulation in the cytoplasm of N4-KO cells as compared to parental Hela cells. Data represent the mean+/−SD of triplicate experiments. **(H)** WB showing activation of Caspase 1 and IL-1β in N4-KO cells. **(I)** Long range PCR assay showing increased mtDNA deletions in spermatocytes from NEURL4+/− mice (N4-het) compared to WT littermates. **(J)** Decreased sperm count in NEURL4 heterozygotes mice compared to WT littermates. Mean of sperm count across 5 random fields for n=4 mice/genotype +/− SD. Statistical significance calculated by two-tailed Student’s t test. * indicates p value <0.05, ** pvalue <0.01, *** pvalue <0.001.

Compared to the nucleus, mitochondria have a simplified DNA damage repair response that relies on DNA base excision repair/single-strand breaks repair (BER/SSBR) activity to contrast oxidative damage (van Houten et al., 2016; Prakash and Doublié, 2015). PARylation of the step limiting enzyme for mtBER resolution, the mitochondrial form of Ligase III (mtLIG3), was previously proposed as a regulatory strategy for mtDNA damage repair (Rossi et al., 2009). Thus, we asked whether mtLIG3 was a target of NEURL4. Indeed, the PARylation mtLIG3, as assessed by IP/WB, was found drastically reduced in both NEURL4-deficient cell lines (**Fig. 3E**) and upon treatment with the generic inhibitor BGP-15 2HCl, but not with PARP1 inhibitor Olaparib (**Fig. 3F lane 2 and 3**). This confirms that NEURL4, rather than previously investigated ARTs, is responsible for the PARylation of mtLIG3. Moreover, in absence of NEURL4 we observed increased accumulation of cytosolic mtDNA (**Fig. 3G**) and corresponding activation of IL-1β processing and caspase activation (**Fig. 3H**), as expected in the case of progressive accumulation of deleterious mutations and mtDNA deletions in N4-KO cells ultimately leading to organelle dysfunction and mtDNA release in the cytosol, where it is recognized as danger-associated molecular pattern (DAMP) by innate immune signaling pathways (Grazioli and Pugin, 2018).

Intriguingly, large-scale mtDNA deletions are widely regarded as a genetic biomarker for susceptibility to male infertility(Jiang et al., 2017; Karimian and Babaei, 2020; Kumar and Sangeetha, 2009). As this provided a possible mechanism for the difficulties we had encountered in the process of generating a whole body NEURL4-KO mouse, we hypothesized that NEURL4-dependent regulation of mtDNA integrity is essential for maintaining male fertility. In support of this hypothesis, we observed that mtDNA derived from the sperm of NEURL4 male heterozygous mice showed an abnormally high rate of deletions (**Fig. 3I**). NEURL4 heterozygous mice also presented reduced sperm count (**Fig. 3J**) and increased sperm agglutination (**Sup. Media A and B**), two gold-standard markers of male infertility. Collectively, these results indicate that NEURL4 is required for the maintenance of mtDNA integrity both *in vitro* and *in vivo,* and suggest this may be at least partially mediated by ADP-ribosylation of DNA damage repair factors like mtLIG3, in a fashion similar to that of PARP1/ARTD1 in the nucleus.

Based on the fact that previous studies had characterized a multitude of mitochondrial ADP-ribosylated proteins across different functions and that mitochondria-associated PARylation is almost completely lost in N4-KO cells (see Figure 2C), we then considered that NEURL4 may be broadly required for mediating the PARylation of variety of mitochondrial proteins. To characterize the role of NEURL4 enzymatic activity in the regulation of mitochondrial functions, beyond the modulation of mtLIG3, we profiled the poly-ADP-ribosylation status of mitochondrial proteins in NEURL4 wild type and KO cells via mass spectrometry-based quantitative proteomics using Stable Isotope Labeling (SILAC). Putative targets were first enriched via binding to the WWE PAR-binding domain of Rnf146 (Darosa et al., 2015) and then identified by LC-MS/MS. With this approach, we identified ~170 putative targets defined as proteins with decreased poly-ADP-ribosylation in both KO cell lines and confirmed in two separate experiments (**Supplemental Fig. S1A**)(**Supplemental Table 1**). In agreement with the intra-mitochondrial localization reported above, NEURL4 putative targets are enriched for proteins located in the matrix/inner mitochondrial membrane (**Fig. 4A**). Moreover, overlay of our results with previously reported total PARylome data (Martello et al., 2016), indicates that several of the mitochondria proteins identified as ADP-ribosylated are in fact included among putative NEURL4 targets (**Supplemental Table 1**). Remarkably, among these targets are factors involved in the regulation of major mitochondrial functions (**Fig. 4B**), including not only DNA damage repair, but also carbon, fatty acid and amino acid metabolism. As expected in presence of putative alterations in the functions of multiple factors/enzymes across diverse mitochondrial pathways, NEURL4-deleted cells present striking morphological and metabolic changes when compared to the parental line. Mitochondria in N4-KO cells are fragmented and present a donut-like shape that has been associated with loss of Δψm in uncoupled mitochondria (Ding et al., 2012; Liu and Hajnóczky, 2011)(**Fig. 4C**). In accord with these phenotypic changes, assessment of mitochondrial bioenergetics by Seahorse respirometry showed a significant reduction in FCCP-induced OCR, with both basal and maximal respiration being severely impaired in N4-KO cells in comparison to the parental cell line (**Fig. 4D**). Moreover, transcriptomic profiling of N4-KO cells by RNA-seq revealed significant reprogramming of gene expression, with activation of gene programs important for metabolic adaptation to mitochondrial dysfunction (**Fig. 4E, Supplemental Fig. S1B and Supplemental Table 2**). Thus, together, the phenotypic and genomic characterization of NEURL4-deficient cells confirms its relevance as a major mitochondrial ART enzyme and indicate that ADP-ribosylation of mitochondrial proteins by NEURL4 is critical for maintaining mitochondria and cellular homeostasis.

**Figure 4.**
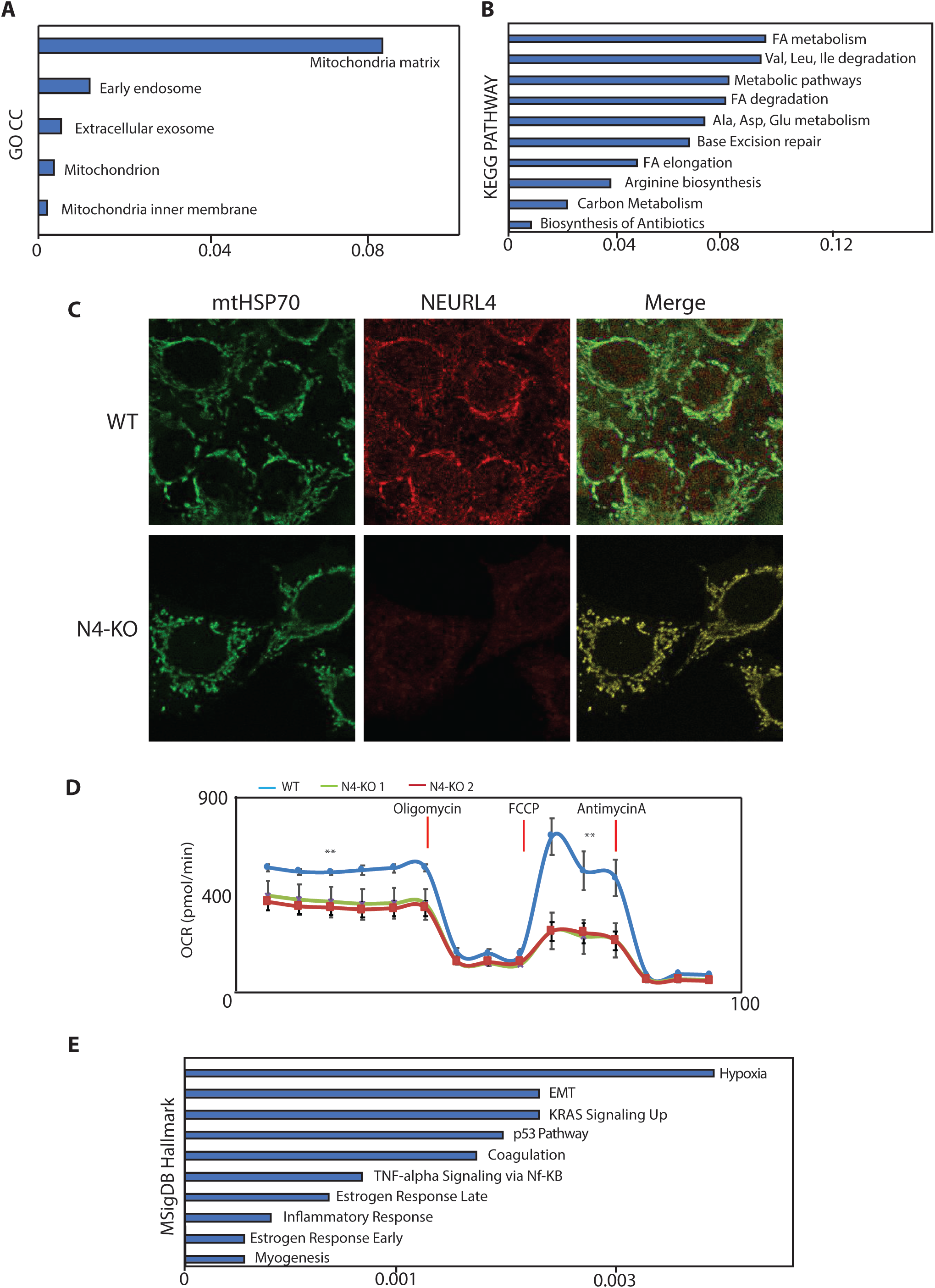
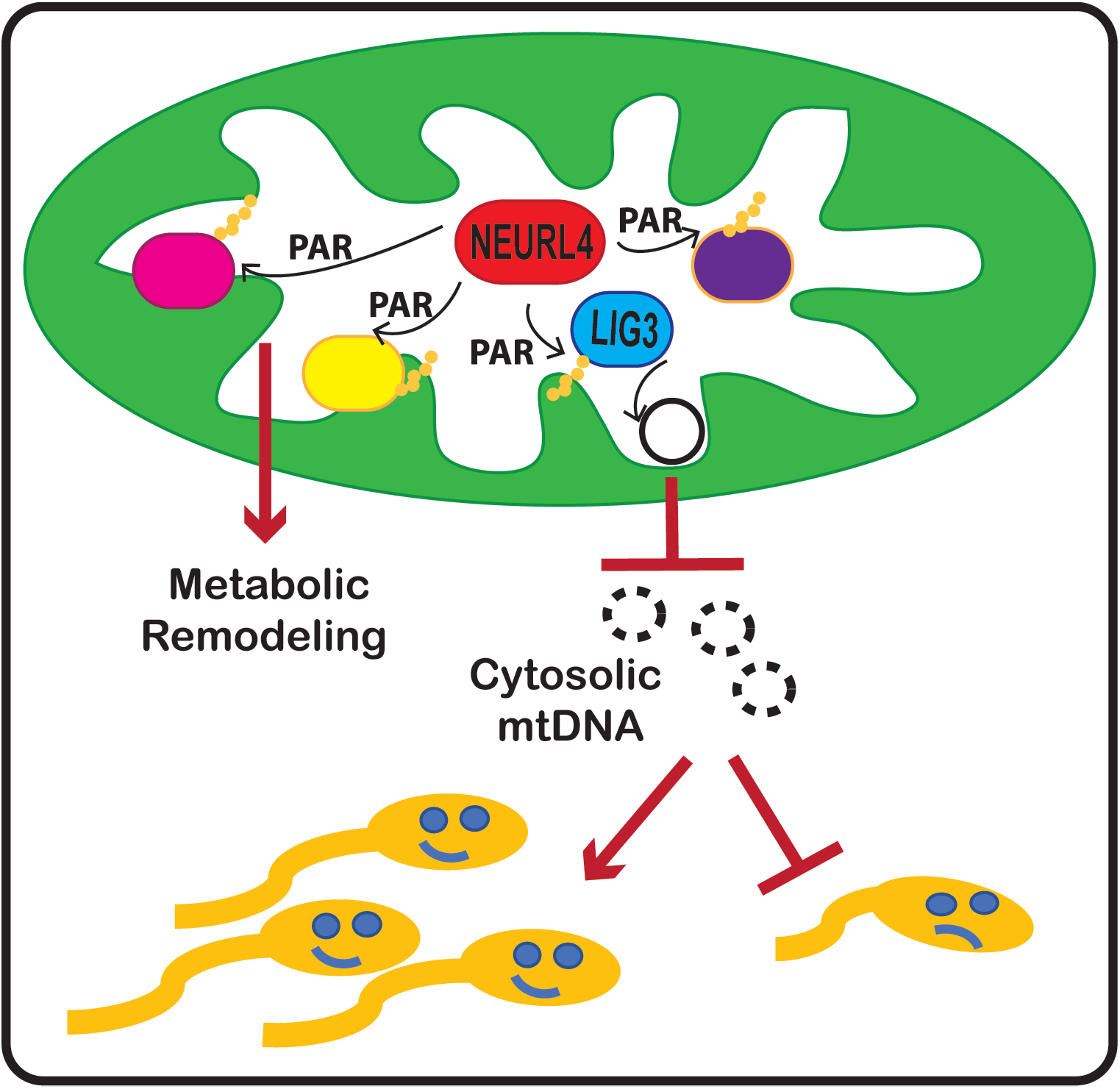
NEURL4-mediated regulation of mitochondrial proteins by PARylation. **(A**) and **(B)** SILAC-based enrichment of PARylated proteins in the absence of exogenous genotoxic stress. Most significant GO terms associated with proteins with decreased PARylation in absence of NEURL4 **(A)**. Most significant pathways associated with decreased PARylation in absence of NEURL4 **(B)**. GO terms and KEGG Pathways enrichment is based on combined score between p-value and Z-score (DAVID). **(C)** Immunostaining showing mitochondrial network changes in Hela cells in absence of NEURL4. **(D)** Decreased basal respiration and impaired FCCP response in Hela N4-KO cells by Seahorse. **(E)** Most significant gene sets from the Molecular Signature Database (MSigDB) enriched among DEGs between WT and N4-KO Hela cells.

## Discussion

As mitochondria represent an ideal environment for post-translational regulation via ADP-ribosylation due to the elevated NAD^+^ levels, it is not surprising that a multitude of mitochondrial proteins were found included among poly-ADP-ribosylated targets identified by large scale proteomic studies (Brunyanszki et al., 2016; Vivelo et al., 2017). However, the ART enzyme/enzymes responsible to mediate their modifications have remained elusive for a long time (Brunyanszki et al., 2016). Previous studies reported the presence of two enzymes with ART activity, SIRT4 and ARTD1, in the mitochondria, but their impact on the mitochodrial PARylome is unclear, with Glutamate dehydrogenase (GDH) being the sole mitochondrial target of SIRT4 identified to date (Herrero-Yraola et al., 2001; Rossi et al., 2009). Here, we have characterized NEURL4 as a novel member of the ARTD family (to be named ARTD-17)(Hottiger et al., 2010) and the long-sought ART enzyme responsible of carrying out ADP-ribosylation in the mitochondria. Our results confirm the enzymatic activity of a predicted ARTD-like domain located at the C-terminus of NEURL4, characterize NEURL4 presence in the mitochondria matrix, where it interacts with mtDNA, and define the NEURL4-dependent mitochondrial PARylome.

Overall, our findings are indicative of a predominant role for NEURL4 enzymatic activity in the regulation of mitochondrial functions. NEURL4 deletion in fact leads to the almost complete loss of PAR synthesis in the mitochondria, in both human and mouse cells. This effect was recapitulated by treating cells with the generic PARPs inhibitor BGP-15 2HCl (Racz et al., 2002) but not with the specific PARP1 inhibitor Olaparib(Menear et al., 2008). Accordingly, profiling of NEURL4-mediated PARylome by mass spectrometry has revealed putative targets implicated in several mitochondrial processes, including metabolites and ions transport, protein synthesis, mitochondrial metabolism, oxidative respiration and mtDNA repair. Even though further studies are needed to elucidate the effect of NEURL4-dependent PARylation on the regulation of each specific target, based on the broad range of mitochondrial functions that are potentially regulated by NEURL4 through ADP-ribosylation, we expected NEURL4-KO cells to present with a severe mitochondrial phenotype. Indeed, in absence of NEURL4 mitochondria appear stressed, with loss of mitochondrial membrane potential and impaired mtDNA integrity. These morphological and functional changes are opposite to those induced by the depletion of ARTD1(Módis et al., 2012; Niere et al., 2008), thus reaffirming that the observed phenotypes are specific to NEURL4 deletion. Among NEURL4 putative targets we identified mitochondrial DNA ligase III (mtLIG3), a key enzyme in DNA replication and repair pathways. Despite the previously reported interaction between ARTD1 and mtLIG3 on mtDNA(Rossi et al., 2009), our data indicate that the enzymatic activity of NEURL4, not ARTD1, is required for basal PARylation of mtLIG3. Further studies will be required to investigate whether there is any crosstalk between the two enzymes and to dissect the molecular mechanisms of mtLIG3 regulation through PAR. Nonetheless, our data indicate that loss of mtLIG3 PARylation, in absence of NEURL4, correlates with decreased mtDNA stability, elevated mtDNA levels in the cytosol and corresponding activation of an innate immune response leading to increased IL-1β processing. Together, these observations are indicative of a key role for NEURL4 in the maintenance of mtDNA integrity, possibly through post-translational regulation of mtLIG3 activity in the base excision repair (BER) pathway. NEURL4 relevance for the maintenance of mtDNA integrity was confirmed *in vivo* as the spermatocytes of male NEURL4^+/-^ heterozygous mice show significant accumulation of mtDNA deletions, which leads to reduced sperm count and increased agglutination. Remarkably, de novo mutations in *Neurl4* gene were recently identified as causative of human male infertility (Hodžić et al., 2020), in agreement with extensive literature linking mtDNA deletions with infertility in both mice and humans(Jiang et al., 2017; Karimian and Babaei, 2020; Kumar and Sangeetha, 2009). On the contrary, ARTD1-KO mice show no major phenotypic differences when compared to wild type if not under genotoxic stress (Niere et al., 2008), again highlighting the pivotal role of NEURL4 in the maintenance of mitochondrial homeostasis under physiological conditions.

## AKNOWLEDGEMENTS

We are grateful to past and present members of the Perissi Lab and Dr. Marc Liesa (UCLA) for feedback and stimulating discussions, and to Dr. L. Aravind for sharing useful insights about the predicted enzymatic activity of NEURL4. We are thankful for the excellent assistance provided by the BUSM Microarray and Sequencing Core Facility (Dr. Yuriy Alekseyev); the BUSM Confocal Microscopy Facility (Dr. Vickery Trinkaus-Randall); the BUSM Analytical Instrumentation Core (Dr. Lynn Deng); the BNORC Adipose Biology and Metabolism Core (Dr. Stephen Farmer) and Mouse Transgenic Core (Dr. Brad Lowell); and the Harvard Electron Microscopy Facility (Dr. Maria Ericsson).

---

**Figure S1 (Supplement to Figure 4).**
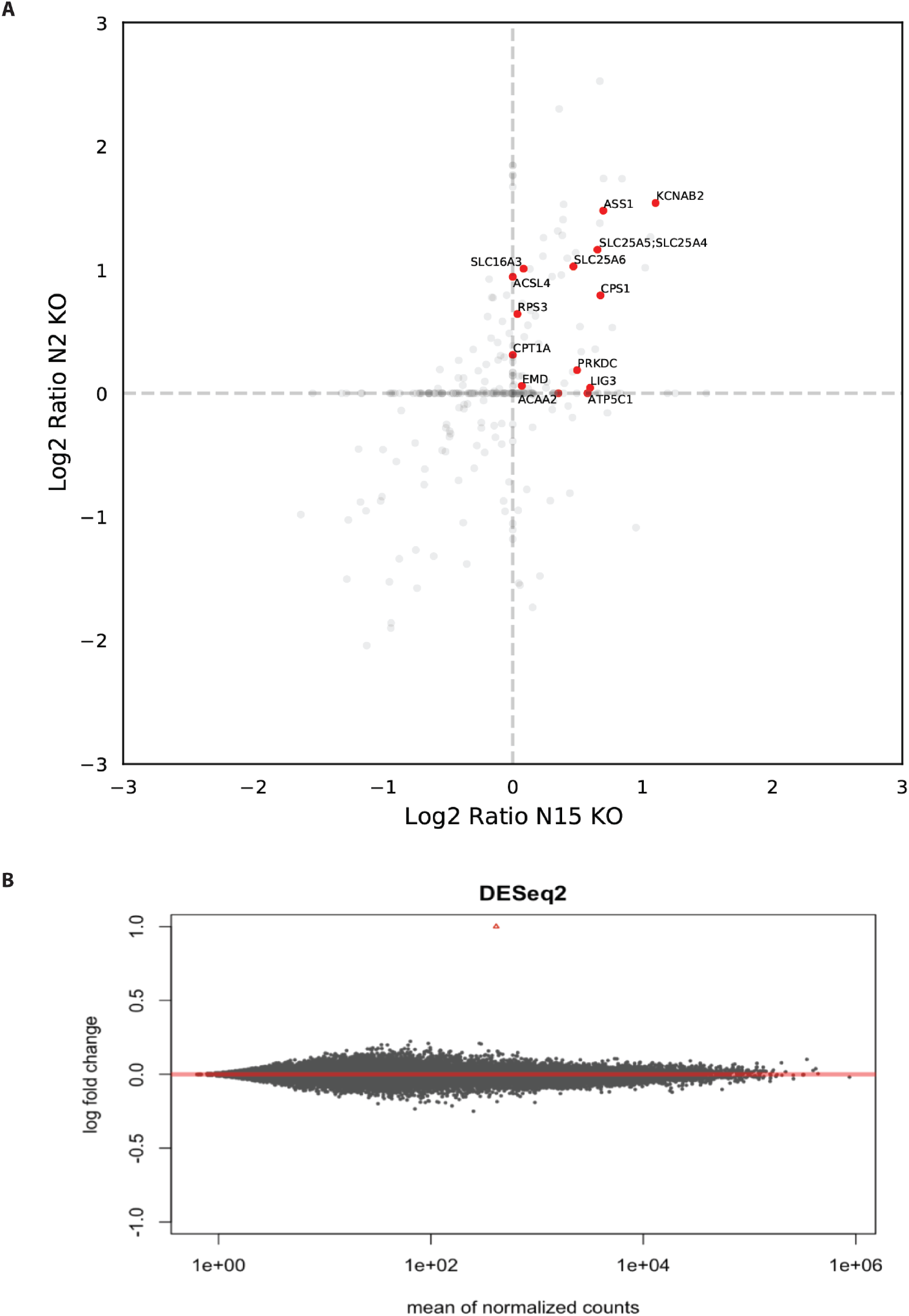
**(A)** SILAC-based enrichment of PARylated proteins identified in WT non-stressed Hela cells using either NEURL4 KO1 or KO2 cells as a background. **(B)** Profiling of DEGs identified in NEURL4 KO1 and KO2 cells as compared to WT parental Hela cells by RNA-seq.

