## Supplementary material for "ADP-ribosylation of mitochondrial proteins is mediated by Neuralized-like protein 4 (NEURL4)": Methods

### STAR*METHODS

| REAGENT or RESOURCE | SOURCE | IDENTIFIER |
|---|---|---|
| Antibodies | | |
| NEURL4 rabbit polyclonal ct | custom | aa 1492-1508 |
| HDAC2 rabbit polyclonal | Santa Cruz | H-54 sc7899 |
| TOM20 mouse monoclonal | Santa Cruz | F-10 |
| mtHSP70 mouse monoclonal | Thermo Scientific | JG1 |
| beta-Tubulin mouse monoclonal | Sigma | D66 |
| ATP5B mouse monoclonal | Molecular Probes | A21351 |
| FLAG-HRP | Sigma | A8592 |
| Poly(ADP-ribose) rabbit | Enzo Life Science | ALX-210-890A Discontinued |
| Cleaved-IL-1β rabbit | Cell Signaling | 83186 |
| LIG3 mouse monoclonal | Santa Cruz | sc-390922 |
| Caspase 1 mouse monoclonal | Santa Cruz | sc-398715 |
| NLRP3 rabbit | Abcam | ab214185 |
| Tot-K-Ac rabbit | Abcam | Ab80178 |
| anti rabbit Rhodamine RedX | Jackson ImmunoReserch | 711-295-152 |
| anti mouse Fluorescein | Jackson ImmunoReserch | 715-095-151 |
| Bacterial and Virus Strain | | |
| Home-made E. Coli | This Paper | N/A |
| Chemicals, Peptides, and Recombinant Proteines | | |
| Lipofectamine 2000 | Invitrogen | 11668019 |
| FCCP | TOCRIS | 04-531-1900 |
| Oligomycin A | Sigma | 75351 |
| Antimycin | Sigma | A8674 |
| DMEM | Gibco | 11965 |
| Olaparib | Cell Signaling | 93852 |
| BGP-15 | Tocris | 6703 |
| FBS | Gibco | 10438 |
| Proteinase K | Sigma | P2308 |
| SUPERasein | Invitrogen | AM2694 |
| Trypsin-EDTA | Sigma | 59418C |
| Protein A-Sepharose | Invitrogen | 101041 |
| Poly(A) Polymerase | NEB | M0276 |
| Phusion® High-Fidelity DNA Polymerases | NEB | M0530 |
| Long Amp Hot Start Taq DNA Polymerases | NEB | M0534S |
| BSA | Fisher | BP9706 |
| Critical commercial Assay | | |
| Quick Start™ Bradford Protein Assay | Biorad | 5000201 |
| ECL | Biorad | 1705060 |
| Faster SYBER Green Master Mix | AB | 4385612 |
| QuickExtract DNA | Epicenter | QE09050 |
| Rneasy Plus | Qiagen | 74136 |
| QuickExtract DNA | Epicenter | QE09050 |
| Experimental Models: Cell Lines | | |
| Hela | ATCC | CCL2 |
| Experimental Models: Organisms/Strains | | |
| NEURL4-het | This paper | N/A |

| | | |
|---|---|---|
| Oligonucleotides | | |
| gRNA NEURL4-exon 15 Fw | Sigma | CACCGACATAGTCACCTTTACCCGG |
| gRNA NEURL4-exon 15 Rv | Sigma | AAACCCGGGTAAAGGTGACTATGTC |
| gRNA NEURL4-exon 2 Fw | Sigma | CACCGCTTGCCATAAAGGTCCACGA |
| gRNA NEURL4-exon 2 Rv | Sigma | AAACTCGTGGACCTTTATGGCAAGC |
| TFAM Fw human | Sigma | TGCTTGGAAAACCAAAAAGACCTC |
| TFAM Rv human | Sigma | TGAATCACCCTTAGCTTCTTGGAA |
| mt-Nd1FW human | Sigma | CAACCCCCTGGTCAACCTCA |
| mt-Nd1Rv human | Sigma | GCCGATCAGGGCGTAGTTTG |
| mt-Cytb divergent Fw human | Sigma | tgaggccaaatatcattctgaggggc |
| mt- Cytb divergent Rv human | Sigma | tttcatcatgcggagatgttggatgg |
| TFAM Fw mouse | Sigma | CTGCACTCTGCCCATCCAAA |
| TFAM Rv mouse | Sigma | CTGAGCATTCGCAGGCCTTT |
| mt-Nd1FW mouse | Sigma | GCCACCTTACAAATAAGCGCTCTC |
| mt-Nd1Rv mouse | Sigma | ACGCAATTTCCTGGCTCTGC |
| mt-Cytb divergent Fw mouse | Sigma | gacgtaaattacggtgacta |
| mt- Cytb divergent Rv mouse | Sigma | tagtcacccggtaatttacgtc |
| Recombinant DNA | | |
| GST-WWE | | N/A |
| NEURL4 FL-FLAG | This paper | N/A |
| NEURL4 DeltaCt-FLAG | This paper | N/A |
| NEURL4 Ct-FLAG | This paper | N/A |
| Software and Algoritms | | |
| GO/pathway enrichment | Huang da et al. 2009 | DAVID |
| GO/pathway enrichment | Kuleshov et al. 2016 | enrichR |

### CONTACT FOR REAGENT AND RESOURCE SHARING

### EXPERIMENTAL MODEL AND SUBJECT DETAILS

HeLa cell were grown in 10% FBS/DMEM supplemented with 0.1 mM MEM non-essential aminoacids, 2mM L-glutamine, 1mM sodium pyruvate. For cells transfection, Lipofectamine 2000 was used following the manufacturer's protocol (Invitrogen). 1 µM of Olaparib (Cell Signaling), 10µg/ml BGP-15 and 10µM Carbonyl cyanide-*p*-trifluoromethoxyphenylhydrazone (FCCP) TOCRIS were used for cell treatments. Two NEURL4 knockout cell lines (N4-KO1 and N4-KO2) were generated using standard CRISPR-Cas9 genome editing technology with two independent sgRNAs (Ran et al., 2013). NEURL4 heterozygote mice were generated by the Boston Nutrition Obesity

Research Center (BNORC) Transgenic Core via injection of ES NEURL4 knockout cells purchased from the KOMP repository. Mice were maintained on standard laboratory chow diet in temperature-controlled facility on a 12-hour light/dark cycle. All animal studies were approved by the Boston University Institutional Animal Care and Use Committee (IACUC) and performed in strict accordance of NIH guidelines for animal care.

### METHOD DETAILS

#### Cell staining

Immunostaining was performed following standard protocols on cells fixed in 4% paraformaldehyde/PBS using Rhodamine RedX (RRX)-conjugated secondary antibodies anti-rabbit and Fluorescein (FITC)-conjugated secondary antibodies anti-mouse (Jackson ImmunoResearch).

#### Protein extraction, Subcellular fractionation, Submitochondrial Localization, *In Vitro* PARylation assay and Immunoprecipitation.

For whole cell extracts preparation, cells were rinsed in PBS, harvested and incubated for 20' on ice in IPH buffer (50 mM Tris-HCl pH 8.0, 150 mM NaCl, 5 mM EDTA, 0.5% NP-40, 50 mM NaF, 2 mM $Na_2VO_3$, 1mM PMSF and protease inhibitor mix). For cytoplasmatic, mitochondrial and nuclear extracts fractionation cells were rinsed in PBS, harvested and re-suspended in gradient buffer (10 mM HEPES pH 7.9, 1mM EDTA, 210 mM Mannitol, 70mM Sucrose, 10mM NEM, 50 mM NaF, 2 mM $Na_2VO_3$, 1mM PMSF and protease inhibitors cocktail) then homogenized via 10 passages through 25G syringe followed by low-speed centrifugation for 10 min. The nuclear pellet was incubated for 20 min in high-salt buffer (10 mM Hepes pH 7.9, 20% glycerol, 420 mM NaCl, 1.5 mM $MgCl_2$, 0.2mM EDTA, 0.5mM DTT, 10mM NEM, 50 mM NaF, 2 mM $Na_2VO_3$, 1mM PMSF and protease inhibitor mix) while the supernatant was recovered and subjected to high-speed centrifugation to separate the mitochondrial pellet from the cytoplasmic fraction. The mitochondrial pellet was incubated for 15 min in lysis buffer (50 mM Tris/HCl pH 8, 300 mM NaCl, 1mM EDTA, 1% Triton X-100, 10mM NEM, 50 mM NaF, 2 mM $Na_2VO_3$, 1mM PMSF and protease inhibitor mix). Blotting with markers of the different fractions (HDAC2 for nuclear extract, mtHSP70 for mitochondrial extracts) was used to assess purity. To examine submitochondrial localization, the isolated mitochondria fraction was treated with either Proteinase K (2, 5 or 10 ng) in ice for 30 min or 50 □g/ml trypsin in ice for 30 min under either isotonic or hypotonic condition, reaction was terminated adding respectively 1mM PMSF or 10% TCA. Concentration of protein extracts was measured using the colorimetric BIORAD assay. Extracts were boiled in SDS sample buffer and loaded 10% Mini-PROTEAN TGX gels (Biorad), prior to transfer onto PVDF membranes (Millipore) and western blotting following standard protocols. For *in vitro* PARylation assay cells were transfected with FLAG-NEURL4-DCt or FLAG-NEURL4-Ct then mitochondrial were isolated and protein extract were endogenous subjected to immunoprecipitation as previously described (Cardamone et al., 2012) to isolate endogenous NEURL4 or FLAG-NEURL4-DCt and FLAG-NEURL4-Ct. Immuno-precipitated proteins were then incubated with $NAD^+$ in reaction buffer (50 mM Tris-Hcl, pH 7.4, 2 mM MgCl2) with or without 2.5 µg of ssDNA (Sigma) at 37 °C for 30 min.

**ChIP assay**

Chromatin immunoprecipitation (ChIP) was performed as described (Perissi et al., 2004). Briefly, approximately $10^7$ cells were cross-linked with 1% formaldehyde at room temperature (~25 °C) for 10 min and neutralized with 0.125 M glycine. For BAT experiments, four biological replicated ChIPs were pooled for sequencing, for both male and female mice. Tissue was minced with curved scissors (slurry like) and put in 10ml tubes containing PBS 1% formaldehyde for crosslink for 15 min at RT and neutralized with 0.125 M glycine. Tissue was then shredded in lysis buffer using a Bullet Blender homogenizer at max power for 5 min 4°C. After sonication, chromatin was incubated with 2µg of antibody at 4 °C overnight. Immunoprecipitated complexes were collected using Sepharose A beads (Life Technologies). After extensive washes the DNA was extracted and purified by phenol/chloroform extraction. ChIP experiments were repeated at least three times and representative results are shown as samples mean between technical replicates +/- standard deviation. Significance was calculated by paired student's T-test.

**RNA Isolation, RT-PCR Analysis and RNA-seq**

RNA was isolated from cell or mice tissue following the manufacturer protocol for the RNeasy Kit (QIAGEN). For the RNA-seq, cells were subjected to standard RNA isolation prior to RNA library preparation following Illumina's RNA-Seq Sample Preparation Protocol. Resulting cDNA libraries were sequenced on the Illumina's HiSeq 2000.

**Immunogold-Electron Microscopy**

For preparation of cryosections cells were rinsed in PBS, harvested and fixed in 4% paraformaldehyde/PBS for two hours then cells were infiltrated with 2.3M sucrose/PBS/glycin 0.2M) and frozen in liquid nitrogen. Frozen samples were sectioned at -120$^{0}$C and transferred to formvar-carbon coated copper grids. Gold-labeling was carried out following standard protocols (Harvard EM Core).

**Mitochondrial content and mtDNA cytoplasmic content**

Total DNA was extracted from cells using QuickExtract DNA Extraction Solution 1.0 (Epicenter) following manufacturer's instructions. To isolate DNA from the cytoplasm cells pellet were resuspended in 500 mL buffer composed of 150 mM NaCl, 50 mM HEPES (pH 7.4), and 25 mg/ml digitonin. The homogenates were mixed for 10 min at room temperature to allow plasma membrane permeabilization, then centrifuged at 1000 g for 10 min to pellet intact cells. Cytosol-containing supernatant was transferred to a new tube and centrifuged at 17,000 g for 10 min to remove any remaining cellular debris. DNA was then isolated from this fraction using the DNA purification kit from Qiagen. Quantitative PCR was performed on whole-cell extracts and cytosolic fractions using mitochondrial-encoded NADH dehydrogenase 1 (mt-ND1) relative to nuclear TFAM was used to determine mitochondrial DNA copy numbers.

**Respirometry**

Cells were plated in Seahorse V.7 multi-well culture plates. The next day, media was replaced by running media (XF Seahorse Assay Media supplemented with 5.5mM glucose, 0.5mM pyruvate and 1mM glutamine) and the plate was placed at 37°C for 1 h (no carbon dioxide). Oxygen consumption was measured at 37°C using a Seahorse XF24 Extracellular Flux Analyzer (Seahorse Bioscience, Billerica, MA). Mitochondrial stress test compounds (10 μM oligomycin, 2.5 μM FCCP and 10 μM antimycin A) were injected through ports A, B, and C, respectively, to measure mitochondrial respiration linked to ATP synthesis, leak, maximal respiratory capacity and non-mitochondrial oxygen consumption according to the manufacturer's instructions.

**Recombinant protein production and Immunochemical methods**

Purified GST tagged WWE domain was obtained by expressing the proteins in *E.coli* BL21(DE3) cells, which was induced with 0.5 mM isopropyl 1-thio-β-Dgalactopyranoside for 4 h at 30 °C. The harvested cells were lysed by sonication in lysis buffer (1mM DTT, 1% Triton X-100 in PBS). GST-WWE fusion protein was purified from cell lysates using glutathione-Sepharose (GSH) 4B beads (Qiagen, Valencia, CA) according to the manufacturer's instructions. The GST-WWE-conjugated agarose was stored at 4 °C in PBS supplemented with 30% glycine.

For pull-down of polyADP-ribosylation proteins, mitochondrial proteins were extracted from Hela cells. GST-WWE-conjugated agarose were added to immobilize the poly-ADP-ribosylation proteins, and bound material was washed extensively in high-stringency buffer (50 mM Tris, pH 7.5; 500 mM NaCl; 5 mM EDTA; 1% NP40; 1 mM dithiothreitol (DTT); 0.1% SDS). Proteins were resolved by SDS-APGE and analyzed by immonoblotting.

**SILAC-based mitochondrial polyADP-ribosylation proteins pull-down and in-gel digestion**

For SILAC experiments, Hela cells were grown in medium containing unlabelled L-arginine and L-lysine ($Arg^0$/$Lys^0$) as the light condition, or isotope-labelled variants of L-arginine and L-lysine ($Arg^{10}$/$Lys^8$) as the heavy condition (Ong et al., 2002). Mitochondrial proteins were extracted from SILAC-labelled Hela wild-type and Neurl4-knockout cells as descripted before. An equal amount of proteins from the two SILAC states was mixed and precipitated by GST-WWE-conjugated agarose and incubating at 4°C for 1h. Precipitated proteins were eluted with SDS sample buffer, incubated with DTT(10mM)

for 10min at 100 °C. Proteins were separated by SDS-PAGE and visualized with Coomassie blue stain, and then were digested in-gel by trypsin overnight using standard methods (Jensen et al., 1999). The resulting peptides were desalted using C18 Tips (Thermo Scientific) per the manufacturer's instructions.

**Mass spectrometric analysis**

Peptides were analysed on a Q-Exactive HF mass spectrometer (Thermo Fisher Scientific) equipped with a nanoflow HPLC system (Thermo Fisher Scientific). Peptides were loaded onto a C18 trap column (3 μm, 75 μm × 2 cm, Thermo Fisher Scientific) connected in-line to a C18 analytical column (2 μm, 75 μm × 50 cm, Thermo EasySpray) using the Thermo EasyLC 1200 system with the column oven set to 55 °C. The nanoflow gradient consisted of buffer A (composed of 2% (v/v) ACN with 0.1% formic acid) and buffer B (consisting of 80% (v/v) ACN with 0.1% formic acid). For protein analysis, nLC was performed for 180 min at a flow rate of 250 nL/min, with a gradient of 2-8% B for 5 min, followed by a 8-20% B for 96 min, a 20-35% gradient for 56min, and a 35-98% B gradient for 3 min, 98% buffer B for 3 min, 100-0% gradient of B for 3 min, and finishing with 5% B for 14 min. Peptides were directly ionized using a nanospray ion source into a Q-Exactive HF mass spectrometer (Thermo Fisher Scientific).

The QE-HF was run using data dependent MS2 scan mode, with the top 10 most intense ions acquired per profile mode full-scan precursor mass spectrum subject to HCD fragmentation. Full MS spectra were collected at a resolution of 120,000 with an AGC of 3e6 or maximum injection time of 60 ms and a scan range of 350 to 1650 m/z, while the MS2 scans were performed at 45,000 resolution, with an ion-packet setting of 2e4 for AGC, maximum injection time of 90 ms, and using 33% NCE. Source ionization parameters were optimized with the spray voltage at 2.1 kV, transfer temperature at 275 °C. Dynamic exclusion was set to 40 seconds.

**Data analysis**

All acquired MS/MS spectra were searched against the Uniprot human complete proteome FASTA database downloaded on 2018_10_26, using the MaxQuant software (Version 1.6.7.0) that integrates the Andromeda search engine. Enzyme specificity was set to trypsin and up to two missed cleavages were allowed. Cysteine carbamidomethylation was specified as a fixed modification. Methionine oxidation, N-terminal acetylation, and ADP-ribosylation on a wide range of amino acid residues (C, D, E, H, K, R, S, T, and Y) were included as variable modifications. Peptide precursor ions were searched with a maximum mass deviation of 6 ppm and fragment ions with a maximum mass deviation of 20 ppm. Peptide and protein identifications were filtered at 1% FDR using the target-decoy database search strategy. Proteins that could not be differentiated based on MS/MS spectra alone were grouped to protein groups (default MaxQuant settings).

**Statistical analysis**

All of the data shown in the histograms are the results of at least three independent experiments and are presented as the mean +/- SEM. Hand ChIPs data are representative of three independent experiments, bar graphs represent the sample mean of three technical replicates +/- SD. The differences between groups were compared using Student's t test. Imaging results and western blot are representative of three independent experiments.
